# The FDA-approved drugs ticlopidine, sertaconazole, and dexlansoprazole can cause morphological changes in *C. elegans*

**DOI:** 10.1101/2020.04.09.034421

**Authors:** Kyle F Galford, Antony M Jose

## Abstract

Urgent need for treatments limit studies of therapeutic drugs before approval by regulatory agencies. Analyses of drugs after approval can therefore improve our understanding of their mechanism of action and enable better therapies. We screened a library of 1443 Food and Drug Administration (FDA)-approved drugs using a simple assay in the nematode *C. elegans* and found three compounds that caused morphological changes. While the anticoagulant ticlopidine and the antifungal sertaconazole caused morphologically distinct pharyngeal defects upon acute exposure, the proton-pump inhibitor dexlansoprazole caused molting defects and required exposure during larval development. Such easily detectable defects in a powerful genetic model system advocate the continued exploration of current medicines using a variety of model organisms to better understand drugs already prescribed to millions of patients.

## 1. Introduction

Imperfect knowledge about biology makes it difficult to anticipate all the effects of a drug and makes empirical testing necessary. The extent of testing needed, however, is often unclear. Famously, the sedative thalidomide taken by pregnant women for morning sickness caused birth defects in more than 10,000 babies [1,2] possibly through the degradation of a transcription factor required for limb development [3,4] - a mechanism that has taken more than 50 years to elucidate. Ideally, a drug would be tested for efficacy and side effects under many circumstances during the lifetime of individuals and even their descendants. Such comprehensive testing, however, is impractical because of pressing needs for medicines to alleviate suffering. Adequate risk assessment of a drug before approval can also be stymied by latency or underreporting of adverse side effects. For example, the cardiovascular risks of using the COX-2-selective nonsteroidal anti-inflammatory drug rofecoxib was only established more than five years after its approval [5]. Aware of such risks with every approved drug, regulatory agencies continue to monitor unanticipated effects after approval (e.g. MedWatch [6] by the Food and Drug Administration (FDA)). This wait-and-watch strategy could be complemented with proactive analyses of the effects of approved drugs in a variety of model systems. With the approval of 59 new drugs by the FDA in 2018 [7], the need for such post-approval studies has heightened.

Most drug testing is done using mice and other mammals under the premise that they have more human-like features. However, such complex animal models can be difficult to work with and are less characterized than simpler animal models that nevertheless capture some fundamental molecular features of human biology. For example, the nematode *C. elegans* has 7,943 genes with human orthologs [8] and uses many conserved mechanisms, including signaling through G-protein coupled receptors that are similar to those most frequently targeted by drugs in humans [9]. In recognition of these features, *C. elegans* has been proposed as a potentially effective model for detecting toxicity of new compounds [10,11]. Such expansion of the set of model systems to test adverse effects of compounds is also supported by the prior inability of popular mammalian models to reveal human toxicity (for example, the harmful effects of thalidomide are detected in rabbits but not in mice [12]). Complementing such toxicity studies with deep analyses on the effects of compounds that have been approved for treating human diseases could be similarly productive. The extensive characterization of *C. elegans* [13] and its short generation time make it ideally suited for testing the effects of a drug under many circumstances throughout the lifetime of individuals and their descendants.

Here we report our serendipitous discovery of obvious morphological changes in the nematode *C. elegans* caused by drugs in current use.

## 2. Materials and Methods

### (a) *C. elegans* culture

Worm strains (N2: wild type, EG6787: *oxSi487 [Pmex-5::mCherry::h2b::tbb-2 3’UTR::gpd-2 operon::gfp::h2b::cye-1 3’UTR] II; unc-119(ed3) III*, AMJ844: *dpy-2(e8) oxSi487; unc-119(ed3)*, AMJ1020: *dpy-2(jam37) oxSi487; unc-119(ed3); bus-8(jam35) X*) were cultured on plates or in liquid at 15°C [14].

### (b) Genome editing

Cas9-based genome editing was used to introduce the *e2698* mutation [15] into *bus-8* (resulting in BUS-8[A131V]) in AMJ844 using sgRNA transcribed *in vitro*, a 60-base single-stranded DNA for homology repair, and the correction of *dpy-2(e8)* into *dpy-2(jam37)* as a co-CRISPR marker [16, 17]. An edited isolate selected based on failure of AluI (New England Biolabs cat. no: R0137L) restriction digests was designated as AMJ1020.

### (c) Levamisole sensitivity assay

Batches of three L4-staged worms from AMJ1020, N2, and EG6787 incubated in 10 µl droplets of varying concentrations of levamisole (tetramisole hydrochloride, Sigma-Aldrich Cat. no.: L9756) in M9 buffer [14] on a coverslip for 60 seconds were scored as paralyzed if they were unable to perform a half-body bend during the next 60 seconds.

### (d) Screen of compound library

AMJ1020 was used to screen a compound library (Selleck Chem Cat. no.: L1300 in dimethyl sulfoxide (DMSO)) for compounds that cause increased fluorescence in the red channel (2 second exposure with 4×4 binning using filter cube: 530 to 560 nm excitation, 570 dichroic, and 590 to 650 nm emission) and/or in the green channel (varying exposures with 4×4 binning using filter cube: 450 to 490 nm excitation, 495 dichroic, and 500 to 550 nm emission). For each compound, 100 µl of worms from ∼12-day liquid culture was aliquoted into each well of 96-well plates along with 3 µl of 10 mM drug solution and imaged ∼48 hours later. Worms were imaged after immobilization with levamisole at a fixed magnification on an AZ100 microscope (Nikon) with a Cool SNAP HQ2 camera (Photometrics) after excitation using a C-HGFI Intensilight Hg Illuminator. Drug-exposed and DMSO-exposed AMJ1020 worms were compared with control EG6787 worms imaged (as in [18]) under the same conditions for each imaging session.

### (e) Secondary assays of positive hits from compound screen

The three drugs identified using the screen were obtained from a second source (Cayman Chemical, cat. no.: 20770 (ticlopidine), 22232 (sertaconazole), 18235 (dexlansoprazole)). Wild-type animals were exposed to newly made solutions of these compounds in DMSO (vehicle) for ∼48 hours, after which L4-staged worms were selected and exposed for an additional ∼24 hours before imaging. To examine morphological defects, live worms were mounted on a 3% agarose pad, exposed for ∼10 minutes in 5 µl of freshly made 3 mM levamisole, and imaged in the red channel without binning (using filter cube: 530 to 560 nm excitation, 570 dichroic, and 590 to 650 nm emission) under non-saturating conditions. Differential Interference Contrast (DIC) imaging of the same animals was performed on a Zeiss Axiovert 200 microscope using a 40x objective. All images being compared were identically adjusted using FIJI (NIH) and/or Illustrator (Adobe) for display.

### (f) Statistical analysis

Fractions of animals showing defects were compared using Wilson’s estimates with continuity correction (method 4 in [19]).

## 3. Results

Bioactive chemicals can have acute effects by disrupting processes that happen in one generation or can have longer lasting effects by disrupting processes that act over many generations. The impact of chemicals on epigenetic processes in particular can be difficult to detect because of the typically transient nature of changes in such processes [20]. We recently reported the stable induction of gene silencing of a fluorescent reporter that lasts for more than 250 generations [18]. Here we combine this reporter of stable transgenerational epigenetic inheritance with mutations that impair the cuticle to generate a drug-sensitive epigenetic sensor (figure 1*a*). Specifically, we generated a *C. elegans* strain in which the expression of two fluorescent proteins – GFP and mCherry – has been silenced epigenetically for many generations [18] and the permeability of the cuticle has been enhanced by mutating the predicted glycosyltransferase *bus-8* [15]. Consistent with previous reports, we found that mutating *bus-8* made worms more susceptible to the paralytic drug levamisole (figure 1*b*).

**Figure 1.**
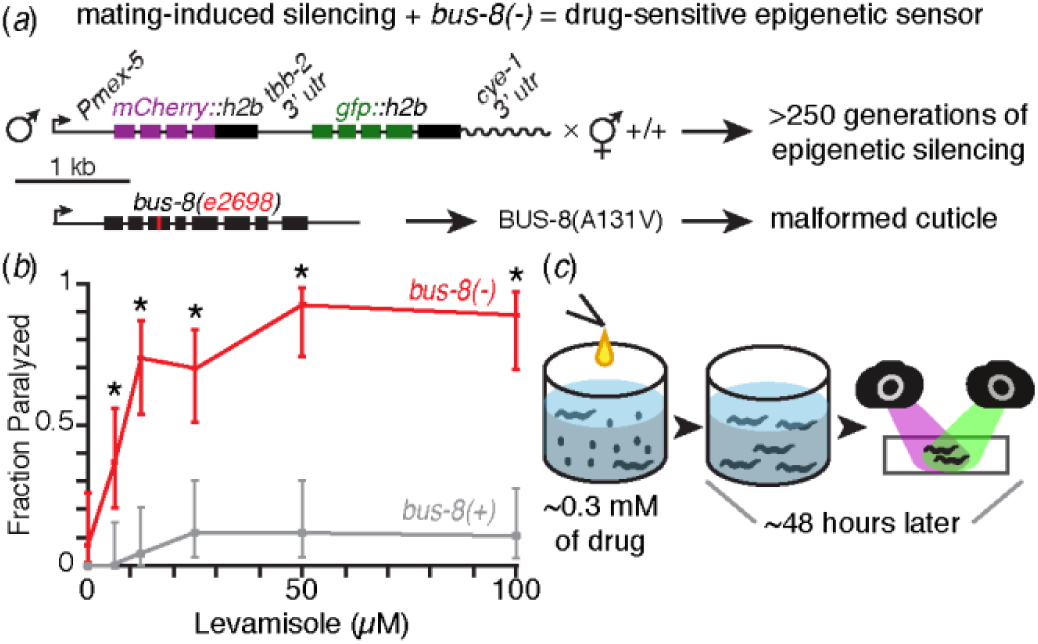
Development and use of a strain for the sensitive detection of bioactive compounds. (*a*) Design of a sensitive sensor. To generate a sensor for drugs that can disrupt physiological and epigenetic processes in *C. elegans*, we combined two features: (i) mating-induced transgenerational gene silencing of a fluorescent reporter (*Pmex-5::mCherry::h2b::tbb-2 3’utr::gpd-2 operon::gfp::h2b::cye-1 3’ utr*) [18] and (ii) the *bus-8(e2698)* mutation that is expected to produce a mutant glycosyltransferase protein BUS-8(A131V), resulting in a malformed cuticle (based on [15]). (*b*) The sensor strain is hypersensitive to the paralytic drug levamisole, consistent with increased permeability to small molecules. Fractions of animals with a mutation in *bus-8* (*bus-8(-)*) and animals without the mutation (*bus-8(+)*) that were paralyzed by different concentrations of levamisole are shown. Error bars indicate 95% CI and asterisks indicate p < 0.05, Wilson’s estimates. (*c*) Schematic of screen for drug-induced changes in fluorescence. Liquid cultures of the sensor strain were exposed to ∼0.3 mM of each drug from a library of 1443 FDA-approved compounds for 48 hours and imaged using fluorescence microscopy in red and green channels.

This drug-sensitive mutant with transgenerationally silenced reporters was grown in liquid culture and exposed to a library of 1443 FDA-approved compounds (figure 1*c*). Two of the tested compounds - nitazoxanide and curcumin - were expected to cause obvious effects based on prior studies, and did so in our screen. Nitazoxanide caused lethality (as reported in [21]), and we additionally detected autofluorescent accumulations in Nitazoxanide-exposed worms that were also visible using bright-field microscopy. Curcumin, which fluoresces under our imaging conditions, caused punctate fluorescence within the intestine, likely due to intra-intestinal accumulation [22,23]. While none of the compounds revived the silenced transgene, three compounds that do not fluoresce as solutions under our imaging conditions caused increased fluorescence in the pharyngeal region: ticlopidine, sertaconazole, and dexlansoprazole. Similar increases were observed in wild-type animals without the silenced transgene and malformed cuticle (figure 2), demonstrating that the standard laboratory strain is susceptible to the effects of these drugs.

**Figure 2.**
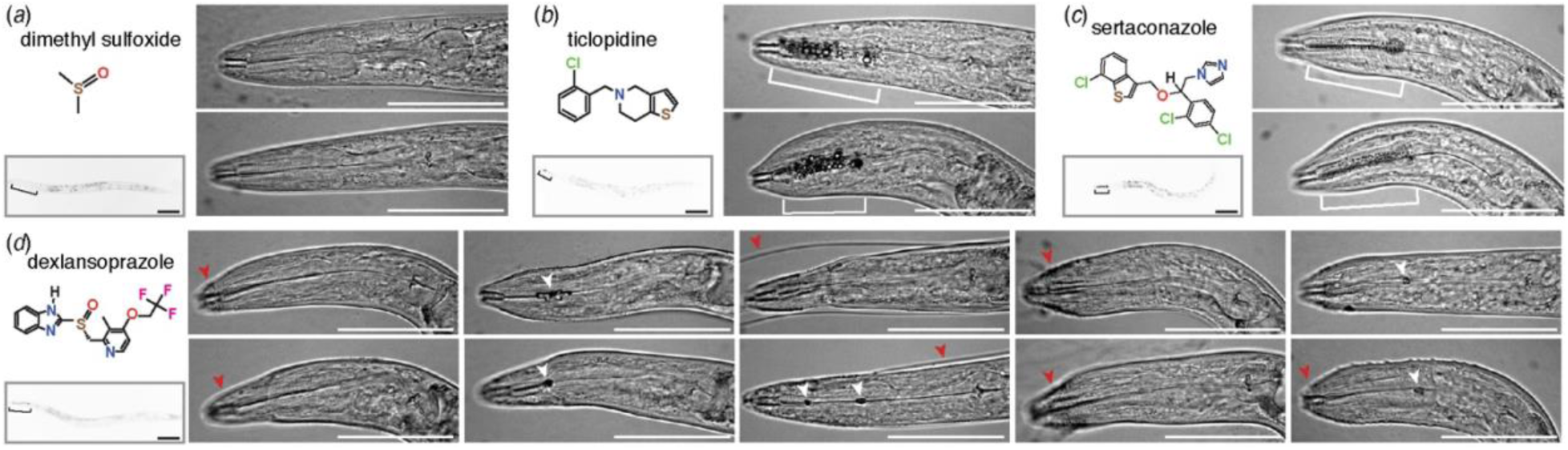
Three FDA-approved drugs can alter the morphology of *C. elegans*. (*a-c*) Ticlopidine and Sertaconazole cause distinct defects in the pharynx upon acute exposure. Unlike the vehicle dimethyl sulfoxide (*a*), ticlopidine (*b*) and sertaconazole (*c*) alter the pharyngeal region in wild-type animals. For each compound, chemical structures (*top left*), induced changes in fluorescence (brackets) at lower magnification in the red channel (*bottom left*; scale bar *=* 100 µm) and induced changes in morphology of the pharynx (brackets and white arrowheads), if any, at higher magnification (*right*; scale bar *=* 50 µm) are shown. Changes in pharyngeal morphology were observed in 0/38 animals upon DMSO exposure, in 25/56 animals upon ticlopidine exposure (p < 0.05, Wilson’s estimates) and in 15/19 animals upon sertaconazole exposure (p < 0.05, Wilson’s estimates). Two different animals are shown for each compound to illustrate the reproducibility of the distinct defects observed in ticlopidine (*b*) and in sertaconazole (*c*). (*d*) Dexlansoprazole causes defects in molting and/or a blockage within the pharyngeal lumen upon exposure during larval development. *Left*, The chemical structure (*top*) and induced changes in fluorescence (brackets) at lower magnification in the red channel (*bottom*; scale bar *=* 100 µm) are shown. *Right*, The defects caused by dexlansoprazole varied from worm to worm. A defect in the pharyngeal region could be discerned under low magnification in 11/82 animals and a selection of 10 animals that exhibit a molting defect (red arrowhead) and/or blockage of the pharyngeal lumen (white arrowhead) is shown.

Each drug caused a distinct defect in the pharyngeal region that was evident even under low magnification using bright-field microscopy. Closer examination of worms exposed to ticlopidine or sertaconazole revealed blebs in the procorpus and metacorpus regions of the pharynx with associated lethality (figure 2*a*-*c*): ticlopidine caused large blebs and 28% lethality (22/78 dead vs 1/39 in DMSO) and sertaconazole caused small blebs and 67% lethality (39/58 dead vs 1/39 in DMSO). These characteristic blebs also occurred upon acute exposure of fourth larval (L4) staged animals to either compound for 48 hours (figure 2*a*-*c*, 45% of ticlopidine-exposed live animals and 79% of sertaconazole-exposed live animals). Both kinds of defects with associated lethality are reminiscent of the recent demonstration of chemical-induced changes in the marginal cells of the pharynx that can concentrate sterols and other hydrophobic compounds [24]. While ticlopidine and sertaconazole were not identified in this study, a wide array of other compounds caused similar defects in the pharynx (Peter J. Roy, personal communication). The reason for the morphological difference between ticlopidine- and sertaconazole-induced changes, however, is unclear and requires further study.

Dexlansoprazole, on the other hand, caused a darkening of the pharynx that was visible under low magnification in L4-staged or young adult animals (11/82 animals). Closer examination of these animals (figure 2*d*) revealed an obstruction in the lumen of the pharynx (6/11 animals) and/or a molting defect (8/11 animals). Unlike the defects caused by ticlopidine or sertaconazole, the defects caused by dexlansoprazole were not associated with lethality (0/38 dead upon acute exposure at the L4 stage) and required exposure during larval development.

Thus, three FDA-approved drugs cause easily detected morphological changes in *C. elegans* that could be analyzed to understand their mechanisms of cellular uptake, modes of action, and spectrum of side effects.

## 4. Discussion

About 4.4 million people in the United States are estimated to have been prescribed ticlopidine, sertaconazole, or dexlansoprazole in a 7-year period (estimate based on those insured between 2007 and 2014 [25]). Ticlopidine is thought to inhibit platelet aggregation by blocking an adenosine-diphosphate receptor [26] and is sometimes used during the placement of stents in coronary arteries. Sertaconazole is thought to inhibit fungal growth by disrupting fungal cell membranes [27] and is used to treat athlete’s foot. Dexlansoprasozle is thought to inhibit gastric acidity by blocking a proton pump [28] and is used to treat gastroesophageal reflux and peptic ulcers. Consistently, these drugs also cause different defects in *C. elegans*, suggesting that different molecular mechanisms could be disrupted by each drug. Intriguingly, the set of genes affected by ticlopidine in human cell lines are negatively connected with genes affected by lansoprazole, a racemic mixture of levolansoprazole and dexlansoprazole (2nd rank on CMap [29]), suggesting a possible convergence of these two disparate compounds on the same molecular effectors. Further analyses using each drug and its variants are necessary to identify active metabolites (if any) and to determine drug-induced molecular changes in *C. elegans* that lead to the observed morphological changes. Similar exploration of approved drugs using a variety of model organisms could open avenues of research that reveal unanticipated effects of accepted medicines.

## Acknowledgements

We thank Edwin Zhang for contributions towards assay development; Pravrutha Raman for help with introducing the *bus-8* mutation; and Norma Andrews, Tom Kocher and members of the Jose lab for comments on the manuscript. This work was supported by the Tier 1 Faculty Incentive Program at the University of Maryland and in part by a grant from the NIH [R01 GM124356].

